# Zinc Regulates the Copper Reductase Activity of Yeast Nucleosomes

**DOI:** 10.1101/2023.09.14.557765

**Authors:** Maria Vogelauer, Chen Cheng, Ansar Karimian, Znala Williams, Kristi Tomlinson, Hooman Golshan Iranpour, Narsis Attar, Siavash K. Kurdistani

## Abstract

Histone H3 is a copper reductase enzyme that generates Cu^1+^ ions for intracellular copper trafficking and metalation of copper-dependent proteins. Whether this activity operates within native nucleosomes, and how it is regulated, has remained unclear. Here, we show that native yeast nucleosomes possess robust copper reductase activity that depends on the histone H3 catalytic site and is strongly influenced by zinc. Zinc acts at two levels, acutely stimulating Cu^2+^ reduction *in vitro* while also establishing a catalytically competent nucleosome state during cell growth. Consistently, impaired zinc uptake lowers cellular Cu^1+^ bioavailability, whereas genetically enhancing nucleosome copper reductase activity partially restores Cu^1+^ availability, improves growth under zinc limitation, and suppresses transcriptional signatures of disrupted zinc homeostasis. These findings reveal the nucleosome as an enzyme complex, identify zinc as a regulator of nucleosome-mediated copper reduction, and support a broader view of chromatin as a metabolic organelle linking metal availability to cellular physiology.

## Introduction

Histones help package eukaryotic DNA and regulate DNA-based processes such as gene expression (1). Yet ancestral histones long predate many of the features with which they are associated in modern eukaryotes. Histone homologs are present in archaea with small genomes, no nucleus, and comparatively limited capacity for epigenetic regulation (2), raising the possibility that histones might therefore possess an older biochemical function that provided a selective advantage before their structural and regulatory roles became elaborated. Motivated in part by the changing metal chemistry that accompanied the emergence of eukaryotic life, we discovered that histone H3 functions as a copper reductase enzyme, catalyzing the reduction of cupric (Cu^2+^) to cuprous (Cu^1+^) ions (3). This reaction has important metabolic consequences because intracellular copper trafficking depends largely on the reduced, cuprous state (Cu^1^⁺) of copper, likely reflecting an evolutionary legacy of early life which arose under low-oxygen, reducing conditions. Copper chaperones such as Atx1 and Ccs1 bind Cu^1+^ and distribute it to pathways that include mitochondrial respiration and antioxidant defense (4). Diminishing histone H3 copper reductase activity through mutation of the critical histidine 113 residue (H3H113) at the putative active site lowers intracellular Cu^1+^ levels and impairs copper-dependent enzymes including cytochrome *c* oxidase (Complex IV of the mitochondrial respiratory chain) and superoxide dismutase 1 (Sod1) (3). These findings identified histone H3 as part of a protein-based mechanism for regulating intracellular copper oxidation state (3).

The metabolic consequences of histone-mediated copper reduction extend beyond copper-dependent enzymes. We previously found that decreasing histone H3 copper reductase activity ameliorates phenotypes associated with frataxin deficiency, the underlying defect in Friedreich’s ataxia (5). Frataxin is required for biosynthesis of iron-sulfur (Fe-S) clusters in mitochondria (6). Fe-S clusters serve as enzymatic or structural co-factors for a wide range of proteins (7). In the absence of frataxin function, the supply of Fe-S clusters declines, adversely affecting many metabolic and informational processes throughout the cell (8,9). Because Cu^1+^ is a major source of Fe-S cluster damage (10,11), diminishing histone H3 copper reductase activity lowered intracellular Cu^1+^ levels and preserved the limited pool of Fe-S clusters in frataxin sufficiently to mitigate the phenotypes associated with their lower rate of production (5). Thus, histone copper reductase activity may contribute to diseases in which Fe-S cluster homeostasis is perturbed (5).

A nucleosome is made up of a tetramer of histones H3 and H4, and two pairs of H2A-H2B dimers, which together wrap 146 bp of DNA (12). The histone H3–H4 tetramer is itself made up of two H3/H4 dimers that associate exclusively through histone H3 residues. The interacting H3-H3’ interface is the proposed catalytic site because that is where Cu^2+^ binds and is likely reduced. Two critical histone H3 residues at the interface, cysteine 110 (C110) and H113, are required for optimal Cu^2+^ binding and reduction *in vitro* (3). Interestingly, *S. cerevisiae* and a subset of other fungi lack the H3C110 residue (13). Our genetic analyses nevertheless suggested that the yeast histone H3 likely retains copper reductase function *in vivo* (3), and recent work showed that reintroducing H3C110 increases histone copper reductase activity and alters copper- and iron-dependent physiology (13) .

Previously, we reconstituted the copper reductase activity of the Xenopus/human histone H3.2–H4 tetramer as well as that of the yeast H3–H4 tetramer containing a reintroduced C110 residue, using recombinant histones *in vitro* (3). In cells, however, the histone H3– H4 tetramer is found primarily in the nucleus and largely in the context of a nucleosome (14), suggesting that histone copper reductase activity may normally operate within chromatin. The nucleosome introduces DNA, additional histones, and posttranslational modifications that could influence access to the catalytic site or modulate its activity. An important unanswered question, therefore, is whether native nucleosomes possess copper reductase activity and, if so, whether that activity is regulated by the cellular environment.

Here, we provide evidence that the native yeast nucleosomes possess copper reductase activity and that activity is influenced by residues at the catalytic site. We adapted a protocol for isolating nucleosomal chromatin from the budding yeast *S. cerevisiae* (15), and developed a modified copper reductase assay to measure its enzymatic activity *in vitro*. Unexpectedly, we found that zinc (Zn^2+^) strongly stimulated nucleosome copper reductase activity *in vitro*, whereas several other divalent metal ions did not. Consistent with these biochemical findings, zinc availability during growth influenced Cu^1+^ production in cells as well as the enzyme activity retained by subsequently purified nucleosomes. Importantly, the gain-of-function H3A110C mutation partially compensated for defects caused by impaired zinc uptake, while growth-promoting effects of H3A110C were strongly dependent on zinc availability. Altogether, our findings suggest that the foundational unit of eukaryotic chromatin in yeast operates as a copper reductase enzyme complex that establishes a functional link between zinc availability, histone catalysis, and intracellular copper redox homeostasis.

## Results

### Purification of native nucleosomes from *Saccharomyces cerevisiae*

To obtain large amounts of native nucleosomes reproducibly, we adapted a protocol developed by Kuznetsov et al. for purification of native chromatin fragments (15). In our protocol (Fig. 1A), yeast cells were grown to logarithmic phase (log phase) and collected. Cell pellets were washed and pre-incubated in a buffer containing the reductant β-mercaptoethanol (βME). Excess βME was subsequently washed off, and the cell wall was digested with recombinant Lyticase in an isotonic buffer to generate spheroplasts. Spheroplasts were then lysed in buffer containing 18% ficoll to disrupt cell membrane while preserving nuclear integrity. Nuclei were separated from cytoplasmic constituents by centrifugation and subjected to Micrococcal Nuclease (MNase) digestion. We empirically optimized the amount of MNase and the duration of digestion to generate mostly mono-, di- and tri-nucleosome chromatin fragments. We then added RNase A to the soluble chromatin to remove contaminating RNA.

**Figure 1.**
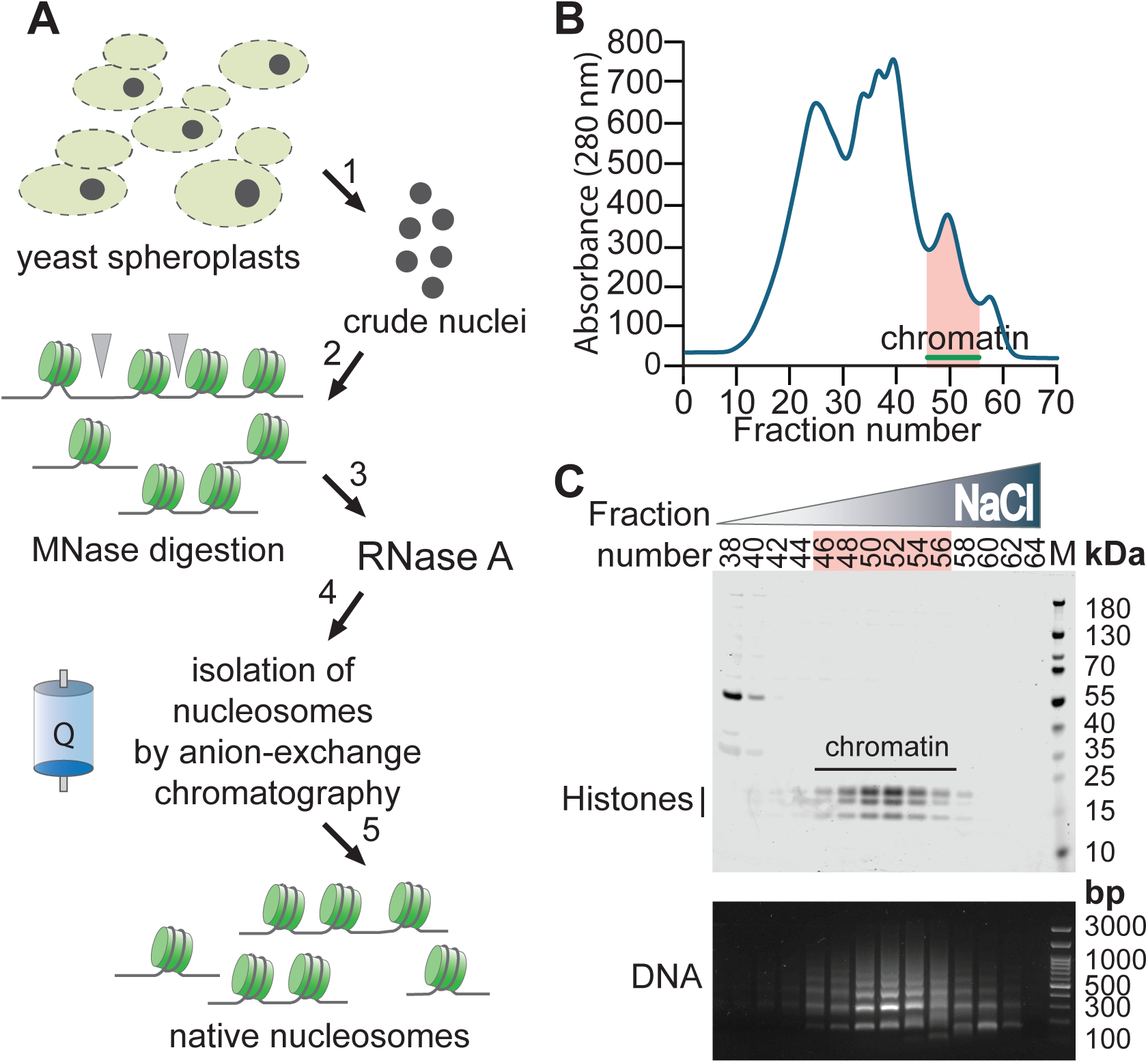
Isolation of native nucleosomes from budding yeast. A, Schematic overview of the protocol for isolating native nucleosomal chromatin from yeast cells. B, Representative chromatogram showing elution of nucleosomal chromatin from an anion-exchange column. C, Representative Coomassie-stained gel of eluted nucleosomal fractions (upper panel) and corresponding nucleosomal DNA isolated from the same fractions (lower panel).

We next purified native nucleosomes using anion-exchange chromatography. Nucleosomes bind to the anion exchange resin primarily through the negatively charged DNA backbone. Bound nucleosomes were then eluted using an increasing salt gradient. Figure 1B displays a representative chromatograph showing the elution profile of chromatin fragments. Gel electrophoresis of collected aliquots identified the fractions containing histones (Fig. 1C upper panel), while analysis of DNA isolated from the same fractions showed that fragments corresponding predominantly to mono-, di-, and trinucleosomes co-eluted with the histones, as expected from a nucleosomal particle purification (Fig. 1C lower panel). A parallel preparation performed without RNase A treatment demonstrated that RNA digestion substantially improved the purity of the nucleosome preparation (Fig. S1A). Fractions containing nucleosomal chromatin were pooled and concentrated for subsequent experiments.

We noticed three bands that co-elute to varying degrees with our nucleosomal fragment preparations (Fig. S1B). To determine their identities and assess whether they might contribute to the measured enzymatic activity, the bands were excised and analyzed by mass spectrometry. The most abundant proteins identified included subunits of the RNA polymerase II, Rpo21 and Rpb2, and potentially Ssa2, an Hsp70-family molecular chaperone associated with the chaperonin-containing T-complex (Fig. S1C). None of these co-eluting proteins has, to our knowledge, been previously associated with copper reductase activity.

For comparisons among assay conditions or between nucleosomes purified from different strains, nucleosome input was carefully normalized using Coomassie-stained gels. Overall, this adapted protocol provides a rapid and reproducible method for isolating native budding yeast nucleosomal chromatin of defined fragment lengths with minimal RNA contamination.

### Copper reductase activity of nucleosomes *in vitro*

We next asked if the isolated nucleosomes reduce Cu^2+^ to Cu^1+^ ions *in vitro*. We adapted an assay that we have previously developed in which bicinchoninic acid (BCA) is used to detect Cu^1+^ production spectrophotometrically. Reactions contained native nucleosomes or control buffer, reduced tris(2-carboxyethyl)phosphine (TCEP) as the electron donor, and BCA in a potassium acetate buffer. Reactions were initiated by addition of Cu^2+^ in the form of Cu^2+^-tricine complex.

In the absence of nucleosomes, Cu^1+^ accumulated slowly over time, whereas addition of wildtype (WT) nucleosomes substantially increased the rate of Cu^1+^ production (Fig. 2A). No significant production of Cu^1+^ occurred in the absence of TCEP, indicating that TCEP, and not the nucleosomes or associated impurities, were the source of electrons (Fig. 2A). The nucleosomes also reduced Cu^2+^ presented in complex with histidine (Fig. S2A). Together, these results indicate that the native budding yeast nucleosomes possess copper reductase activity *in vitro*.

**Figure 2.**
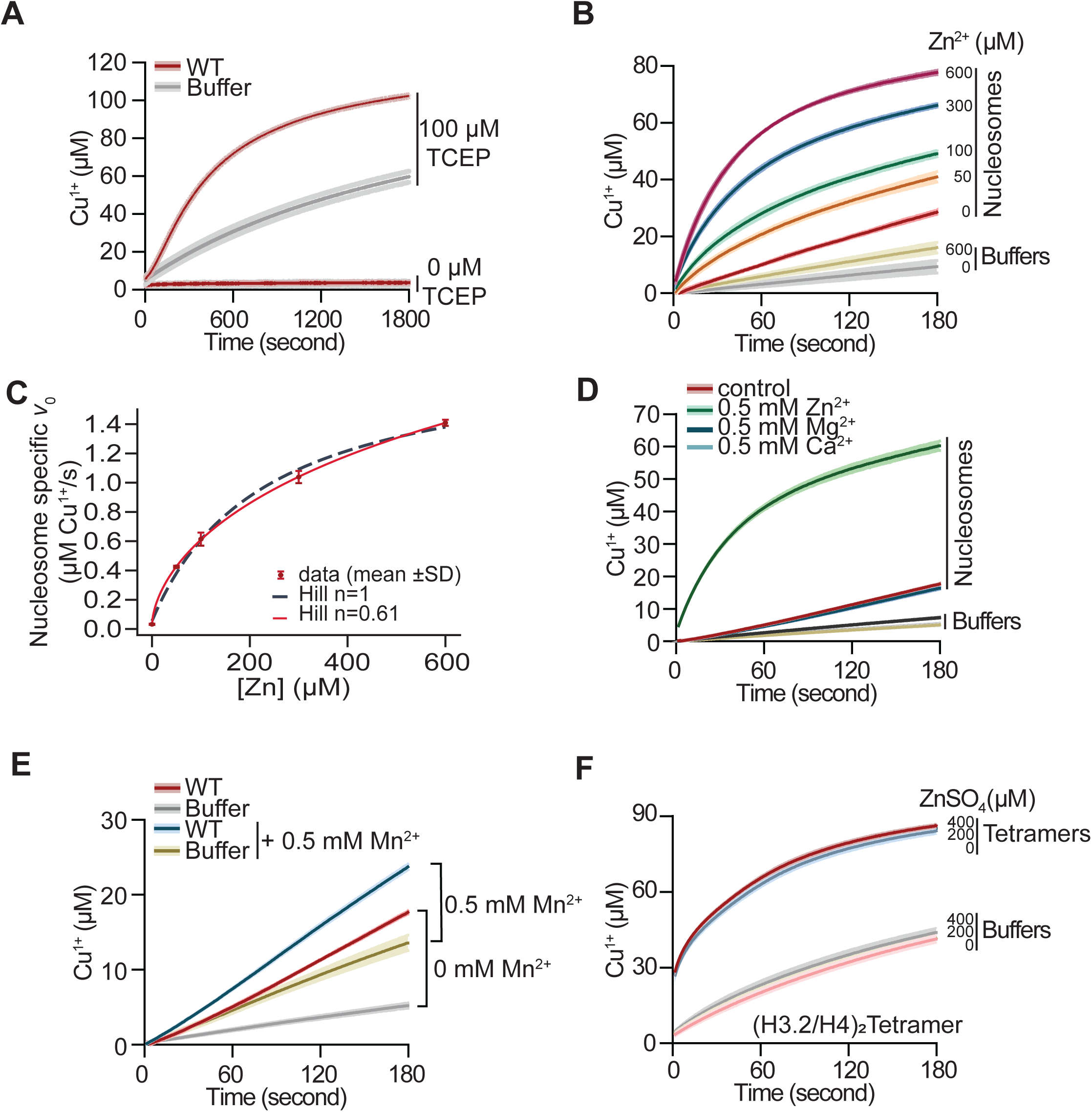
Zinc stimulates the copper reductase activity of native yeast nucleosomes. A, Progress curves of Cu^2+^ reduction by WT nucleosomes or buffer control in the presence or absence of the reducing agent TCEP, using a 1:8 Cu^2+^-tricine complex. Lines and shading represent the mean ± standard deviation (SD) from three assays. B, Progress curves of Cu^2+^ reduction by WT nucleosomes or buffer control using a 1:8 Cu^2+^-tricine complex at pH 7.0 in the presence of the indicated concentrations of ZnSO_4_. C, Nucleosome-specific initial velocities (*v*₀) from the reactions in (B) plotted as a function of Zn^2+^ concentration and fit to Hill models with the Hill coefficient fixed at 1 or allowed to vary. D, Cu^2+^ reduction by WT nucleosomes or buffer control in the presence of the indicated concentrations of Ca^2+^ or Mg^2+^. E, Cu^2+^ reduction by WT nucleosomes or buffer control in the presence or absence of Mn^2+^. F, Progress curves of Cu^2+^ reduction by recombinant H3.2–H4 tetramers or buffer control in the presence of the indicated concentrations of ZnSO_4_.

### Optimal copper reductase activity of native nucleosomes is dependent on zinc ions

Divalent metal ions, including Mg^2+^, Ca^2+^, Mn^2+^, Ni^2+^ and Zn^2+^, can bind to nucleosomes and influence the DNA or nucleosome structure in subtle ways (16,17). Zinc is also a co-factor for many transcription factors and enzymes in eukaryotes (18,19). Interestingly, alterations in zinc levels are associated with changes in copper homeostasis in humans (20). We therefore asked whether divalent metal ions influence nucleosome copper reductase activity *in vitro*.

Addition of Zn^2+^ stimulated the copper reductase activity of WT nucleosomes in a dose-dependent manner (Fig. 2B and S2B). To quantify this effect, we calculated the nucleosome-specific initial velocity (*v*_0_) from the first 15 seconds (s) of the reaction after subtraction of the corresponding control buffer (Fig. S2C). Over the range tested, the copper reduction rate increased approximately 18-fold, from 0.083 µM Cu^1+^/s in the absence of zinc to 1.52 µM Cu^1+^/s at 600 µM zinc. The Zn^2+^ dose-response was better fit by a Hill model in which the Hill coefficient was allowed to vary (n=0.614 ± 0.077) than by a model with n fixed at 1 (Fig. 2C and S2D). A Hill coefficient below 1 is consistent with more than one Zn^2+^ interaction or binding site. This could reflect sites of differing affinity, negative cooperativity, or heterogeneity within the nucleosome population, although these possibilities cannot be distinguished from the dose-response data alone. Thus, zinc strongly enhances the rate of copper reduction by nucleosomes *in vitro* over a broad concentration range.

The stimulatory effect was selective among the divalent metal ions tested. Ca^2+^, Mn^2+^, Mg^2+^ or Ni^2+^ either failed to stimulate nucleosome-specific copper reduction or altered Cu^1+^ production predominantly through nucleosome-independent effects (Fig. 2D, 2E and S2E). Cadmium (Cd^2+^), however, with a valence electron configuration identical to zinc, also stimulated the copper reductase activity of nucleosomes (Fig. S2F), further supporting the selective nature of this stimulation. In addition, zinc had no effect on the copper reductase activity of Xenopus/human histone H3.2–H4 tetramers (Fig. 2F), suggesting that the effects of zinc is specific to nucleosomes.

### Zinc availability during growth establishes nucleosome copper reductase activity

Because Zn^2+^ stimulates nucleosome-mediated Cu^2+^ reduction *in vitro*, we asked whether zinc availability during growth also influences the enzymatic activity of subsequently purified native nucleosomes. WT cells were grown in media containing different concentrations of zinc, and nucleosomes were purified and assayed under identical conditions. Nucleosomes isolated from cells grown in zinc-free medium displayed the lowest copper reductase activity, whereas nucleosomes from cells grown in standard zinc-replete (2.2 µM) or in excess zinc (100 µM) conditions exhibited progressively higher rates of Cu^2+^ reduction (Fig. 3A). Quantification of the initial reaction rates confirmed a zinc-dependent increase in activity with the largest increase occurring between 0 and 2.2 µM zinc (Fig. 3B and S3A). Importantly, nucleosomes isolated from cells grown in 100 µM zinc remained more active than those from cells grown in 2.2 µM zinc even when zinc was omitted from the *in vitro* reaction (Fig. S3B and C), indicating that zinc availability during growth establishes a persistent difference in nucleosome copper reductase activity.

**Figure 3.**
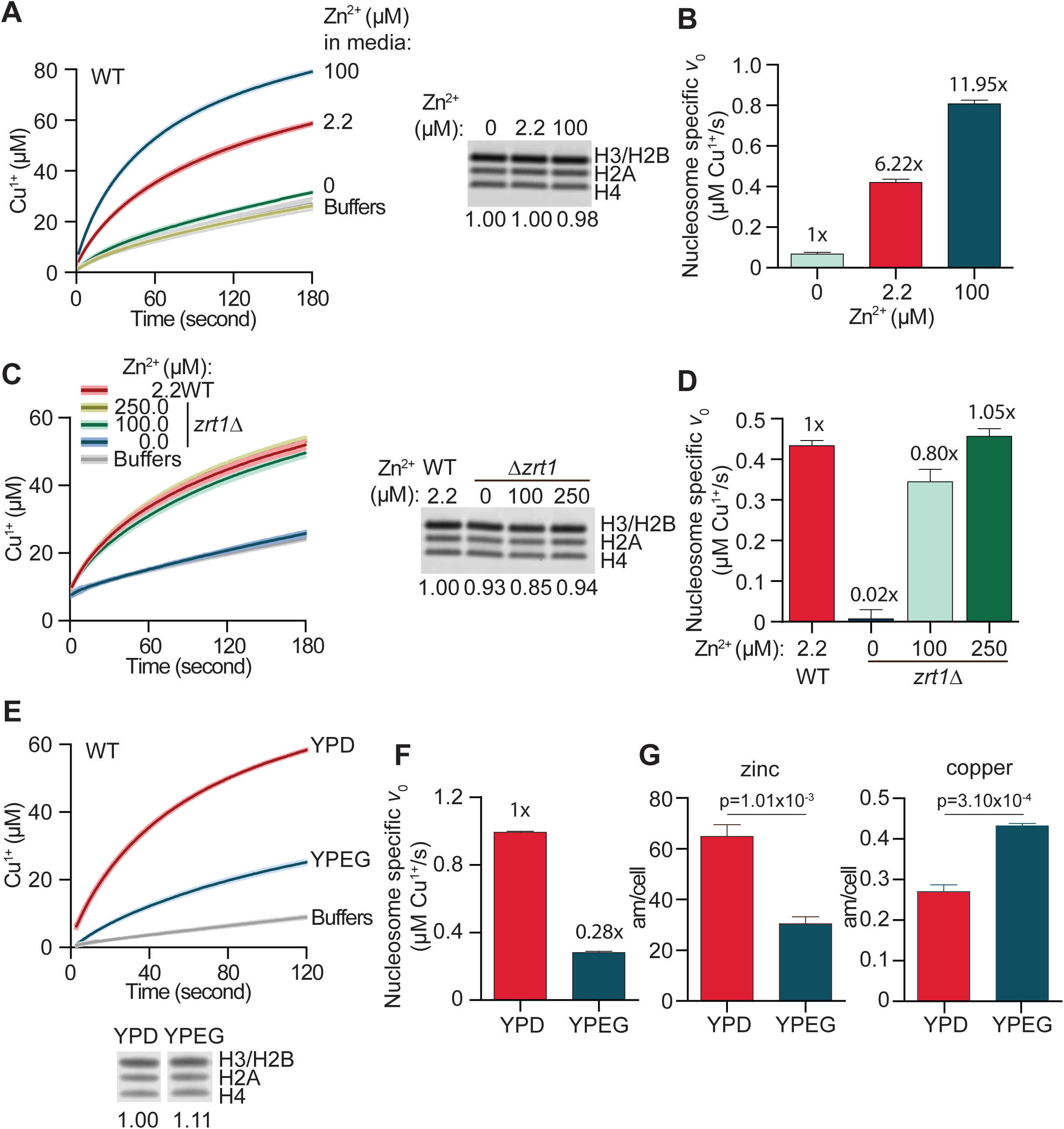
Zinc availability during growth determines the copper reductase activity of purified yeast nucleosomes. A, Left, progress curves of Cu^2+^ reduction by WT nucleosomes purified from cells grown in zinc-free medium or in medium supplemented with 2.2 or 100 µM Zn^2+^. Reactions contained 500 µM ZnSO_4_. Right, Coomassie-stained gel of nucleosomes used in the assay, with relative loading indicated. B, Nucleosome-specific initial velocities (*v₀*) of the reactions shown in (A) with indicated fold-increase relative to nucleosomes isolated from cells grown in zinc-free medium. C, Left, progress curves of Cu^2+^ reduction by nucleosomes purified from WT or *zrt1Δ* cells grown in medium containing the indicated concentrations of Zn^2+^. Reactions contained 500 µM ZnSO_4_. Right, Coomassie-stained gel of nucleosomes used in the assay, with relative loading indicated. D, Nucleosome-specific initial velocities (*v*₀) of the reactions shown in (C), and indicating relative levels to WT nucleosomes from cells grown in 2.2 µM Zn^2+^. E, Progress curves of Cu^2+^ reduction by WT nucleosomes purified from cells grown under fermentative (YPD) or respiratory (YPEG) conditions. Reactions contained 500 µM ZnSO_4_. Coomassie-stained gel of nucleosomes used in the assay, with relative loading indicated is shown below. F, Nucleosome-specific initial velocities (*v*₀) of the reactions shown in (E) with levels relative to YPD indicated. G, Intracellular zinc (left) and copper (right) measured by inductively coupled plasma mass spectrometry (ICP-MS) in cells grown in YPD or YPEG. P values were calculated using Welch’s *t*-test.

To determine whether cellular zinc uptake contributes to this effect, we analyzed nucleosomes isolated from *zrt1Δ* cells lacking the high-affinity zinc transporter (21). In cells grown in zinc-free medium, deletion of *ZRT1* reduced nucleosome copper reductase activity nearly to background levels. Increasing the zinc concentration in the growth medium progressively restored activity, with 250 µM zinc restoring the activity of *zrt1Δ* nucleosomes to approximately that observed for WT cells grown under standard 2.2 µM zinc conditions (Fig. 3C, 3D). These findings further support a role for cellular zinc availability in establishing the copper reductase activity of native nucleosomes.

We next compared nucleosomes isolated from cells grown under fermentative (YPD) or respiratory (YPEG) conditions. Nucleosomes purified from YPEG-grown cells displayed lower copper reductase activity than those isolated from YPD-grown cells (Fig. 3E, 3F). Inductively Coupled Plasma Mass Spectrometry (ICP-MS) analysis revealed that cells grown in YPEG contained less intracellular zinc but more copper than cells grown in YPD (Fig. 3G). In addition, *ZRT1* expression was reduced during respiratory growth (Fig. S3D), providing a potential explanation for the lower cellular zinc abundance under these conditions. Together, these findings indicate that zinc availability during growth establishes, or strongly influences, the copper reductase capacity retained by purified native nucleosomes.

### Mutations at the histone H3 putative catalytic site alter native nucleosome copper reductase activity

A pair of histidine residues at the H3-H3’ (H3H113) interface is important for Cu^2+^ binding and reduction both *in vitro* and in cells (3). Mutation of H3H113 to alanine is lethal in yeast (22,23), a phenotype previously attributed to potential destabilization of the H3-H3’ interface (24). However, introduction of cysteine at histone H3 alanine 110 (H3A110C), which enhances the copper reductase activity of yeast H3 but does not contribute directly to H3-H3’ interactions (12), rescues the lethality of H3H113A (3). This finding suggested that impaired copper reductase activity contributes to the lethality associated with the H3H113A mutation.

Guided by the cancer-associated mutations in H3H113, we previously generated a yeast strain carrying the H3H113N mutation in both copies of histone H3 gene (3). Genetic analyses indicated that H3H113N behaves as a hypomorphic allele and retains sufficient enzyme activity to support viability. Cells carrying the H3H113N mutation (*H3^H113N^*) have lower intracellular Cu^1+^ levels and defects in the activity of copper-dependent enzymes. Consistently, the H3–H4 tetramers bearing the H3H113N mutation displayed lower copper reductase activity *in vitro* (3). Conversely, replacement of yeast H3A110 with cysteine (H3A110C), a residue present in most eukaryotic histone H3 proteins but absent from certain fungi (13), increases the copper reductase activity of yeast H3–H4 tetramers and improves growth under copper-limiting conditions (3).

To determine if the H3H113N and H3A110C mutations similarly affect the copper reductase activity of native nucleosomes, we isolated nucleosomal chromatin fragments from the corresponding yeast strains and assayed copper reduction in the absence or presence of zinc in the reaction (Fig. 4A and B). Nucleosome input was normalized using Coomassie-stained gels (Fig. 4C). In the absence of zinc in the assay, *H3^H113N^* nucleosomes reduced Cu^2+^ less efficiently than WT nucleosomes, whereas the *H3^A110C^* nucleosomes displayed increased enzyme activity (Fig. 4A). Quantification of the maximal reduction rates showed that the *H3^H113N^*and *H3^A110C^* nucleosomes exhibited 73% and 121% of WT activity, respectively (Fig. S4A).

**Figure 4.**
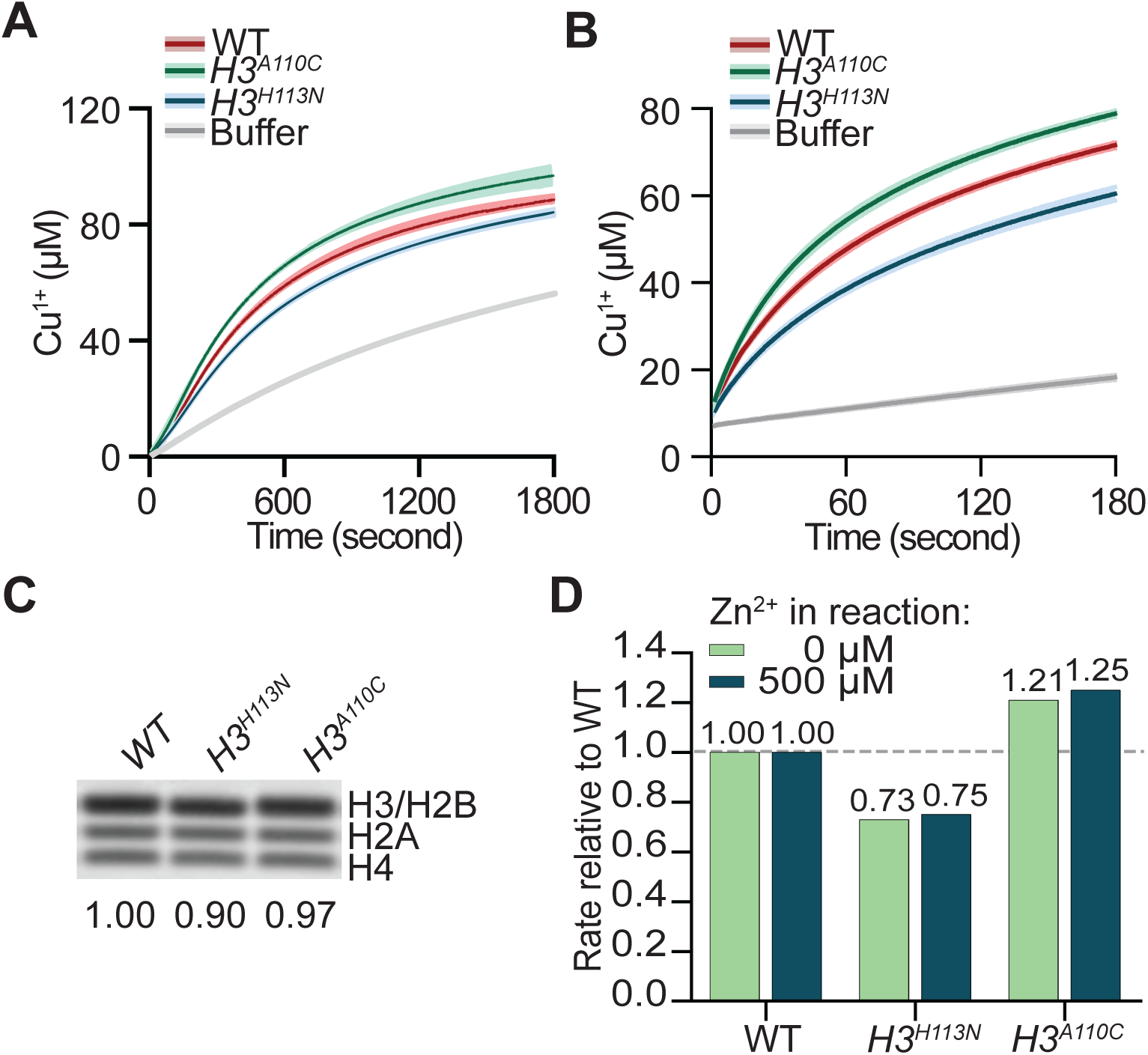
Mutations at the histone H3 interface alter native nucleosome copper reductase activity. A, Progress curves of Cu^2+^ reduction by WT, H3H113N, and H3A110C nucleosomes in reactions without added ZnSO_4_. B, As in (A), but with 500 µM ZnSO_4_ added to the reaction. C, Coomassie-stained gel of nucleosomes used in the assays shown in (A) and (B). D, Copper reduction rates relative to WT for H3H113N and H3A110C nucleosomes in the absence or presence of 500 µM ZnSO_4_. Rates correspond to *V*_max_ for reactions without added zinc and *v*₀ for reactions containing 500 µM ZnSO_4_.

Similar effects were observed when reactions were performed in the presence of 500 µM zinc (Fig. 4B). Measurement of the initial reaction velocities demonstrated reduced activity of *H3^H113N^* nucleosomes and enhanced activity of *H3^A110C^* nucleosomes relative to WT (Fig. S4B). Importantly, the relative effects of both mutations were nearly identical in the presence and absence of zinc in the reaction, suggesting that zinc enhances nucleosome copper reductase activity without altering the intrinsic contributions of key residues at the histone H3 active site (Fig. 4D).

### Zinc availability influences the nucleosome copper reductase activity *in vivo*

To obtain *in vivo* evidence that zinc availability influences the enzymatic activity of nucleosomes, we grew WT and *zrt1Δ* yeast cells under zinc-limiting conditions and assessed intracellular levels of Cu^1+^. To do this, we used a reporter plasmid to read out the activity of the Cup2 transcription factor (3), which is activated directly by Cu^1+^ and does not bind Cu^2+^ (25,26) (Fig. 5A). Importantly, neither Cup2 DNA binding nor its ability to activate transcription depends on zinc (26–28). The reporter contains GFP under the control of the *CUP1* promoter, a major transcriptional target of Cup2 (29,30). We previously showed that the reporter activity responds proportionally to changes in Cu^1+^ levels produced by both physiological and genetic perturbations (3).

**Figure 5.**
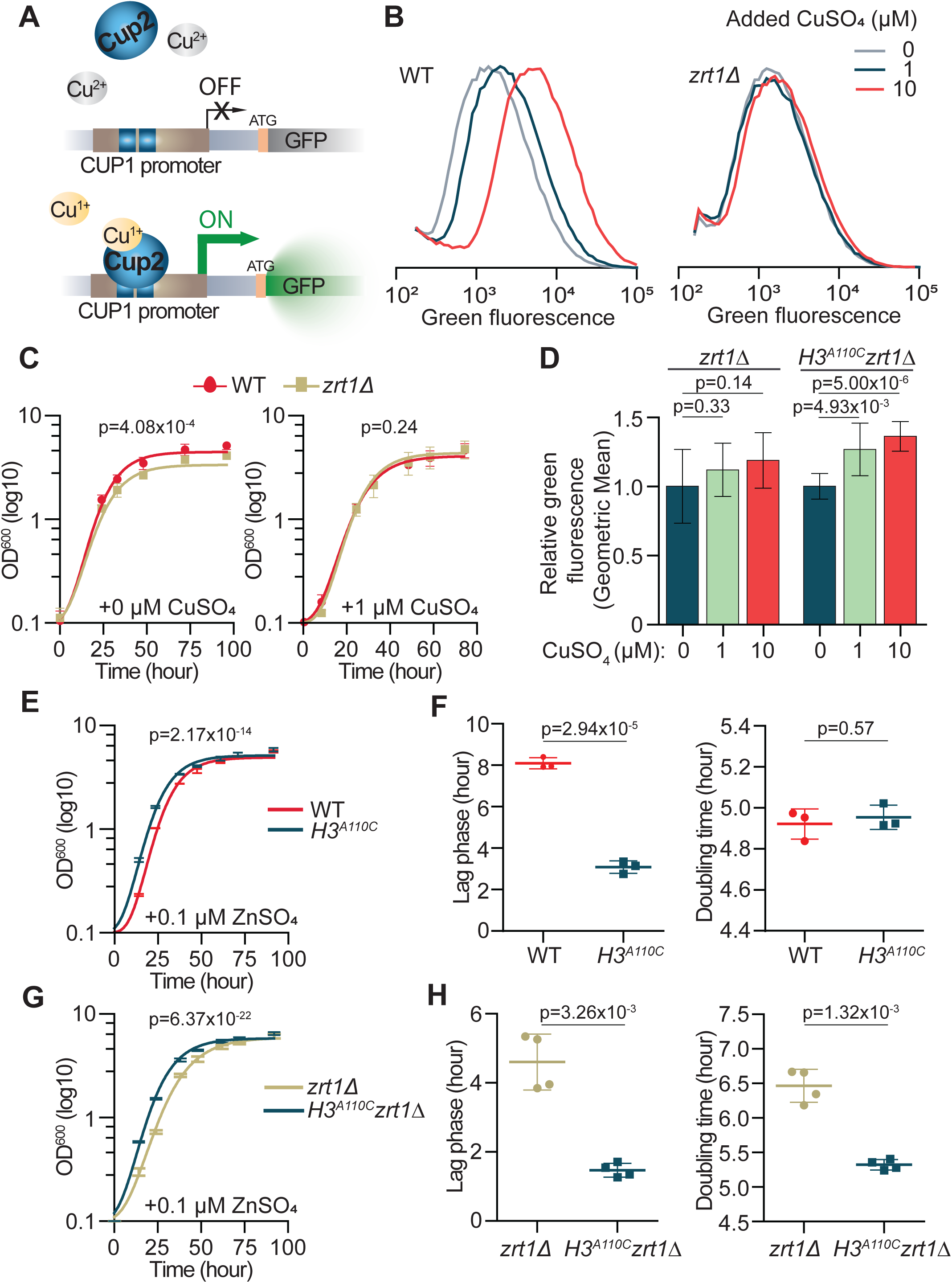
Zinc availability promotes histone H3-dependent Cu^1+^ bioavailability in cells. A, Schematic of the Cup2-dependent pCUP1-GFP reporter used to assess intracellular Cu^1+^ bioavailability. B, Average flow-cytometry distributions of WT or *zrt1Δ* cells carrying the pCUP1-GFP reporter and grown in respiratory SCEG-ura medium containing 0.25 µM ZnSO_4_ and 0, 1, or 10 µM added CuSO_4_. Data are from eight experiments. The basal copper concentration of SCEG-ura medium is approximately 0.16 µM. C, Growth curves of WT and *zrt1Δ* cells in SCEG containing 0.1 µM ZnSO_4_ without or with 1 µM added CuSO_4_. Lines show mean ± SD from three experiments. P values were calculated using an *F*-test comparing the fitted growth curves. D, Relative geometric mean GFP fluorescence of *zrt1Δ* and *H3^A110C^ zrt1Δ* cells carrying the pCUP1-GFP reporter and grown in SCEG-ura containing 0.25 µM ZnSO_4_ and the indicated concentrations of added CuSO_4_. Data are from eight experiments. P values were calculated using Welch’s *t*-test. E, Growth curves of WT and *H3^A110C^* cells in SCEG containing 0.1 µM ZnSO_4_. P values were calculated using an *F*-test comparing the fitted growth curves. F, Lag phase and doubling time calculated from the growth curves shown in (E). P values were calculated using Welch’s *t*-test. G, Growth curves of *zrt1Δ* and *H3^A110C^zrt1Δ* cells in SCEG containing 0.1 µM ZnSO_4_. P values were calculated using an *F*-test comparing the fitted growth curves. H, Lag phase and doubling time calculated from the growth curves shown in (G). P values were calculated using Welch’s *t*-test.

Under both standard fermentative and respiratory growth conditions, GFP expression was similar between WT and *zrt1Δ* yeast cells. However, addition of excess copper to media increased the GFP signal dose-dependently in WT but much less so in *zrt1Δ* yeast cells, indicating that intracellular Cu^1+^ level is diminished when zinc ions are limiting (Fig. 5B and S5A). Because the reporter output depends on Cup2-dependent transcription, these data suggest that Cu^1+^ levels are diminished in nuclei of *zrt1Δ* cells. Consistent with zinc limitation, *zrt1Δ* cells also exhibited impaired growth relative to WT cells under both respiratory and fermentative conditions (Fig. 5C and S5B, left panel). Addition of copper abolished this growth difference (Fig. 5C and S5B, right panel).

We reasoned that increasing the copper reductase activity of nucleosomes might partially compensate for reduced zinc uptake. To test this possibility, we introduced the gain-of-function H3A110C mutation into the *zrt1Δ* background and measured intracellular Cu^1+^ levels using the pCUP1-GFP reporter. The H3A110C mutation consistently increased the GFP signal under respiratory conditions (Fig. 5D), but not under fermentative growth conditions (Fig. S5C), possibly reflecting lower zinc levels in respiratory conditions.

Consistent with these findings, during respiratory growth cells carrying the H3A110C mutation exhibited improved growth under zinc-limiting conditions compared to WT cells (Fig. 5E). Quantification showed that the mutation primarily shortened the lag phase, with little to no effect on the doubling time (Fig. 5F). These effects were not observed in zinc-replete conditions (Fig. S5D). The growth advantage conferred by H3A110C was even more pronounced in the absence of *ZRT1*. In the *zrt1Δ* background, H3A110C shortened the lag phase and also significantly reduced the doubling time (Fig. 5G, 5H), indicating that increasing nucleosome copper reductase activity can partially compensate for impaired zinc uptake. Together, these results support a model in which zinc availability in cells influences intracellular Cu^1+^ levels by modulating the copper reductase activity of nucleosomes.

### Zinc availability is required for the full growth advantage conferred by increased nucleosome copper reductase activity

To further investigate the relationship between zinc availability and nucleosome-mediated copper reduction, we examined the growth of cells lacking the copper-binding metallothionein Cup1. The *cup1F8\** strain, which carries a premature stop codon in every copy of the *CUP1* gene, exhibited impaired growth by potentially altering copper distribution, availability, or toxicity (31,32). We reasoned that increasing nucleosome copper reductase activity through the gain-of-function H3A110C mutation might compensate for this defect.

Under standard zinc-replete conditions (2.2 µM ZnSO_4_), *cup1F8\** cells exhibited a pronounced growth defect characterized by a lag phase that was 92% longer than that of WT cells and a doubling time that was increased by 127% (Fig. 6A-C and 6G, 6H). Introduction of the H3A110C mutation substantially alleviated both defects, reducing the differences relative to WT cells to 13% for lag phase and 29% for doubling time (Fig. 6G, 6H).

**Figure 6.**
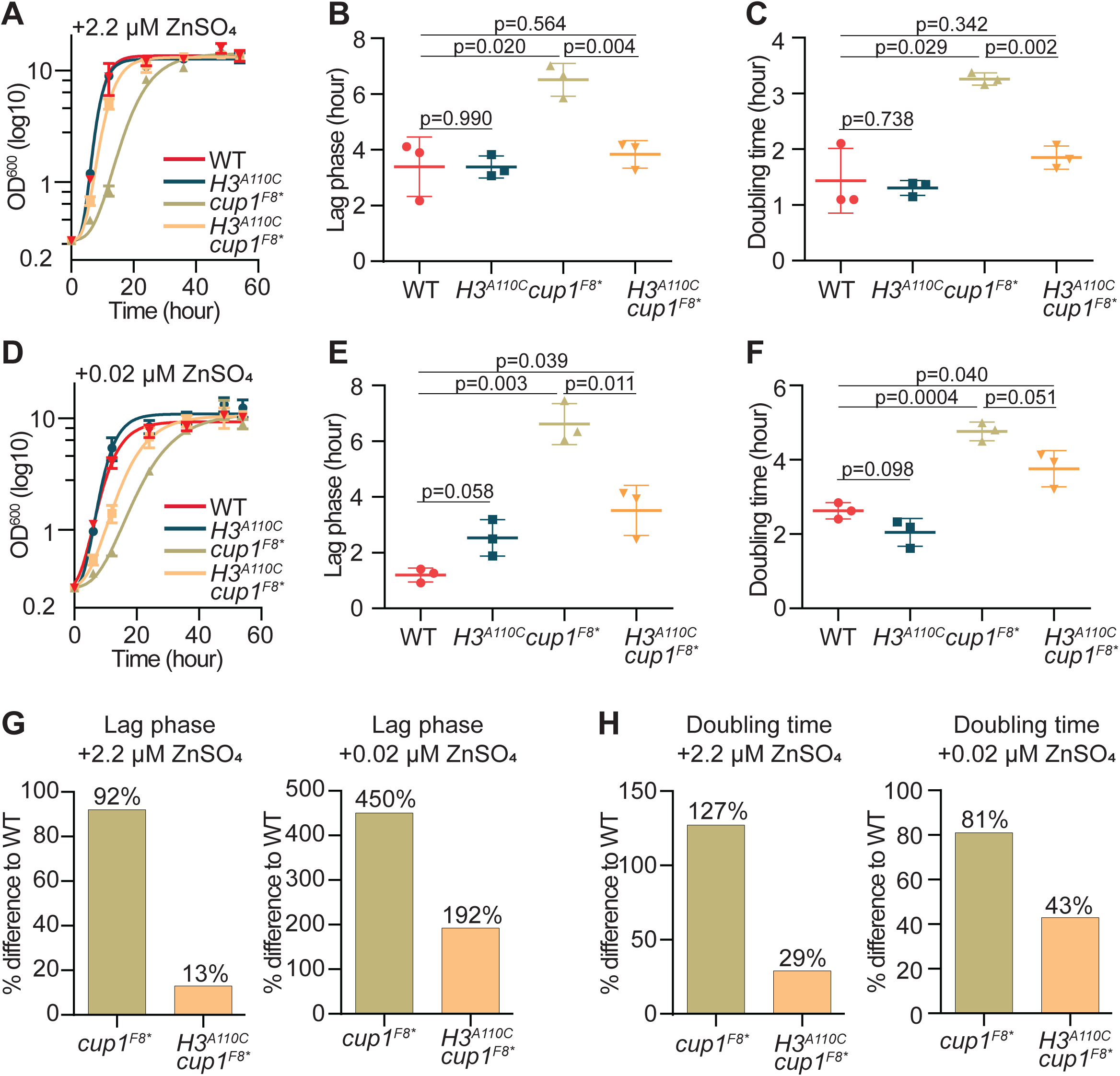
Rescue of the *cup1F8\** growth defect by H3A110C depends on zinc availability. A, Growth curves of the indicated WT and mutant strains in zinc-replete medium containing 2.2 µM ZnSO_4_. B, Lag phase calculated from the growth curves shown in (A). C, Doubling time calculated from the growth curves shown in (A). D, Growth curves of the indicated strains under zinc-limiting conditions containing 0.02 µM ZnSO_4_. E, Lag phase calculated from the growth curves shown in (D). F, Doubling time calculated from the growth curves shown in (D). G, Percentage difference from WT in lag phase for *cup1F8\** and *H3^A110C^cup1F8\** cells under zinc-replete (left) and zinc-limiting (right) conditions. H, Percentage difference from WT in doubling time for *cup1F8\** and *H3^A110C^cup1F8\** cells under zinc-replete (left) and zinc-limiting (right) conditions. P values in (B), (C), (E), and (F) were calculated using Welch’s *t*-test.

In contrast, under zinc-limiting conditions (0.02 µM ZnSO_4_), the growth benefit conferred by H3A110C was markedly diminished (Fig. 6D-F). The lag phase of *cup1F8\** cells increased by 450% relative to WT cells and remained 192% longer even after introduction of the H3A110C mutation (Fig. 6E, 6G). Similarly, the doubling time of *cup1F8\** cells was increased by 81%, and the H3A110C mutation reduced this difference only to 43% (Fig. 6F, 6H).

Consistent with this genetic interaction, mRNA-seq data from Cheng *et al.* (13) showed broad induction of zinc-homeostasis genes in *cup1F8\** cells, including *ZAP1* and genes involved in zinc uptake and intracellular distribution (Fig. S6). Introduction of H3A110C largely restored this expression pattern toward WT, providing additional evidence that increased histone copper reductase activity alleviates the zinc-homeostasis defect associated with loss of Cup1.

Thus, although increasing nucleosome copper reductase activity partially compensates for the loss of Cup1 under both growth conditions, its beneficial effect is substantially greater when zinc is abundant. These findings indicate that the growth advantage conferred by increased nucleosome copper reductase activity is dependent on zinc availability, consistent with the zinc dependence of nucleosome copper reductase activity observed *in vitro*.

## Discussion

We have previously established that the Xenopus laevis (Xl) histone H3–H4 tetramer, which is identical at the protein level to the human H3.2–H4 tetramer, possesses cupric reductase activity *in vitro*. We also provided complementary genetic and molecular evidence that histone H3 functions as a copper reductase in *S. cerevisiae* cells (3). However, the WT yeast H3–H4 tetramer does not appear to assemble efficiently into the correct structure *in vitro*, precluding direct investigation of its potential copper reductase activity. Yeast histone H3 also lacks the H3C110, a residue required for full enzymatic activity of the Xl/human H3–H4 tetramer. Here, we provide evidence that native yeast nucleosomes possess copper reductase activity *in vitro* and that substitutions at the histone H3 catalytic site alter this activity in the directions predicted from previous genetic and biochemical studies. These findings further support the idea that histone H3 functions as an oxidoreductase within nucleosomes and that the fundamental repeating unit of chromatin is itself an enzyme complex. In this sense, chromatin may be considered a “metabolic organelle” that participates directly in cellular metabolism.

A central finding of this study is that zinc influences nucleosome copper reductase activity at two distinct levels. First, Zn^2+^ added directly to the *in vitro* reaction acutely and strongly stimulated Cu^2+^ reduction by native nucleosomes, whereas several other divalent cations had little or no nucleosome-specific stimulatory effect. Second, zinc availability during cell growth altered the catalytic activity retained by nucleosomes after purification. Nucleosomes isolated from cells grown under zinc-replete or zinc-supplemented conditions were more active than those isolated from cells grown in zinc-free medium, and nucleosomes from cells grown in 100 µM zinc remained more active than those from cells grown in 2.2 µM zinc even when no zinc was added to the subsequent *in vitro* assay. We did not determine how much zinc, if any, remained associated with nucleosomes after purification. However, differential zinc carryover alone is unlikely to explain the growth-dependent effect, because addition of Zn^2+^ to the reaction did not restore the low activity of nucleosomes isolated from cells grown in zinc-free medium. Thus, zinc appears to influence nucleosome copper reductase activity both acutely, through its presence during the reaction, and more persistently, by establishing a functional state of the nucleosome during growth.

The effects of substitutions at the histone H3 catalytic site provide an important constraint on how these zinc-dependent effects are interpreted. H3H113N decreased, whereas H3A110C increased, native nucleosome copper reductase activity. Importantly, the relative effects of these substitutions were largely unchanged by the addition of zinc to the *in vitro* reaction. This suggests that acute zinc stimulation does not fundamentally alter the contribution of these residues to catalysis. At the same time, the persistent effect of zinc availability during growth indicates that zinc can influence a property of the nucleosome that cannot simply be recreated by adding zinc to the purified complex.

Several lines of evidence connect this growth-dependent effect of zinc to histone-mediated copper reduction in cells. Loss of the high-affinity zinc transporter *ZRT1* decreased Cu^1+^ bioavailability as measured by the Cup2-dependent reporter. Importantly, the gain-of-function H3A110C substitution, which increases histone H3 copper reductase activity, increased the Cu^1+^ reporter signal under respiratory conditions and partially suppressed the growth defects associated with impaired zinc uptake. Conversely, the ability of H3A110C to improve growth was substantially greater when zinc was replete than when zinc was limiting. These genetic interactions connect zinc availability specifically to a perturbation of the histone copper reductase itself. Together with the biochemical data, these findings support a model in which zinc availability modulates the contribution of chromatin to cellular copper redox homeostasis.

The acute and persistent effects of zinc may arise through at least partially distinct mechanisms. The immediate stimulation observed when Zn^2+^ is added to the reaction could reflect direct interactions with DNA or histones that enhance electron transfer or product release. In contrast, the effect of zinc availability during growth could reflect a more durable change in nucleosome structure, composition, histone modifications, or metal occupancy that survives purification. Although we have not determined whether zinc remains associated with purified nucleosomes, the inability of exogenously added zinc to restore the activity of nucleosomes isolated from zinc-free cells argues that the persistent effect cannot be explained simply by the amount of free zinc present during the enzymatic reaction. The subunitary Hill coefficient observed for acute Zn^2+^ stimulation is consistent with more than one zinc interaction or binding environment and could reflect sites of differing affinity, negative cooperativity, or heterogeneity within the nucleosome population. Zinc binding to DNA has, for example, been proposed to alter the electronic properties of DNA and facilitate charge transfer (33–35). Zinc has also been reported to interact with H3C110 (36), raising the possibility of an interaction near the catalytic site. However, several observations argue against a simple model in which zinc acts solely through H3C110. Yeast histone H3 lacks C110 yet exhibits robust zinc stimulation, and the relative effects of the H3H113N and H3A110C substitutions are largely preserved regardless of whether zinc is added to the reaction. Together, these observations suggest that zinc can influence nucleosome copper reductase activity in at least two ways, through acute stimulation of an already competent nucleosome and through a growth-dependent effect that establishes or maintains that competent state.

The respiratory-growth experiments provide an additional illustration of this relationship. Nucleosomes isolated from cells grown on nonfermentable carbon sources displayed lower copper reductase activity despite the presence of more total cellular copper, whereas cellular zinc abundance and *ZRT1* expression were reduced. Although these observations do not establish a causal pathway, they reinforce the distinction between total cellular copper and the capacity to generate bioavailable Cu^1+^. They also suggest that metabolic state may regulate histone copper reductase activity in part through changes in zinc homeostasis.

The relationship between zinc and copper has long been recognized, although it has generally been considered in terms of metal abundance, transport, competition, or incorporation into metalloproteins. Zinc is required for the function of hundreds of enzymes and regulatory proteins, while proteins such as Sod1 depend on both zinc and copper for full activity (19,37,38). In multicellular organisms, alterations in the copper-to-zinc ratio are associated with oxidative stress, inflammation, aging, and a variety of disease states (20). An especially intriguing parallel comes from the green alga *Chlamydomonas*, in which zinc deficiency causes cellular copper hyperaccumulation while simultaneously producing phenotypes characteristic of copper deficiency (39). Our findings suggest that this apparent paradox may in part reflect zinc-dependent regulation of copper oxidation state. If zinc limitation decreases production of Cu^1+^, cells could accumulate copper while remaining functionally deficient in the reduced copper required for intracellular trafficking and metalation. Whether nucleosome copper reductase activity contributes to the *Chlamydomonas* phenotype remains unknown, but the observation illustrates why copper oxidation state may be an important, and often overlooked, component of zinc-copper homeostasis.

Altogether, our findings establish a previously unrecognized connection among zinc homeostasis, chromatin, and intracellular copper oxidation state. These observations expand the emerging view that histones are not solely structural and regulatory components of the genome but can directly influence cellular metabolism (40–42). The location of this catalytic activity within the highly conserved H3-H3’ interface further raises the possibility that metabolic functions contributed to the selective pressures shaping histone and chromatin evolution, particularly as changing environmental oxygen and metal availability altered the chemical demands placed on early eukaryotic cells. Our findings further suggest that histone mutations or chromatin states that alter copper reductase activity could have metabolic consequences that extend beyond conventional effects on gene regulation. Such effects may be particularly relevant in human diseases in which histones are mutated or copper homeostasis, mitochondrial function, or Fe-S cluster metabolism are perturbed.

## Materials and Methods

### Yeast strains and growth media

*Saccharomyces cerevisiae* strains used in this study are described in Supplementary Table 1. Fermentative and respiratory growth media is 0.17% yeast nitrogen base (Difco and Sunrise Science), 0.5% (NH_4_)_2_SO_4_ with either 2% dextrose (SC) or 2% EtOH/2% Glycerol (SCEG), respectively, and supplemented with amino acids. Amino acid drop out mixes and metal free yeast nitrogen base were purchased from Sunrise Science. Bacteria for histone expression were BL21(DE3) for histone H4 and Lyticase, and BL21(DE3) pLysS for histone H3 and were grown in 2xTY medium (1.6% Bacto Tryptone, 1% Yeast Extract, 0.5% NaCl) supplemented with 100 µg/ml ampicillin or carbenicillin.

### Strain generation for yeast experiments

*ZRT1* was deleted in WT (*OCY1131)* and *H3^A110C^*(NAY616) (3) cells using CRISPR-Cas9 as described in (43). Primers used to create the targeting plasmid are zrt1-pCAS9_F (GGTCTTCCACACATCATCAGGTTTTAGAGCTAGAAATAGC) and pCAS9rev (_p_AAAGTCCCATTCGCCACCCG). The resulting plasmid was then transformed together with the repair-template produced by primer extension of primers zrt1REPf (ACTGCAAGTATAGACAATAAAACAACAGCACAAATATCAAAAAAGGAATTACCAAAGCGA) and zrt1REPr (ATAAAATATGAAATAGAATCTATATGGAACATGCAGAATTTCGCTTTGGTAAAAGGAATT). Successful deletion was verified by PCR.

### Isolation of native nucleosomes from *Saccharomyces cerevisiae*

#### Preparation of crude nuclei using recombinant Lyticase

For isolation of yeast nucleosomes, we adapted the protocol by Kuznetsov et al. (15) with some modifications. 2 liters (L) of yeast cells were grown in indicated media to an OD_600_ of 1.5-2, washed once with deionized water (ddH_2_O), and the wet weight of the pellet was determined. Cell pellets were then flash frozen and stored at -80°C. Cells were thawed and resuspended in 40 ml preincubation solution (60 mM β-mercaptoethanol, 100 mM Tris-HCl pH 8) and incubated for 15 minutes at room temperature under constant agitation. Cells were resuspended in 5 ml/g cells Lyticase buffer (10 mM β-mercaptoethanol, 0.7 M Sorbitol, 0.75% Yeast Extract, 1.5% Peptone, 10 mM Tris-HCl pH 7.5) and incubated with recombinant Lyticase for 30 minutes at 30°C with constant agitation. The amount of Lyticase was established empirically. The degree of spheroplastization was ascertained spectroscopically and microscopically by comparing dilutions in ddH_2_O before and after Lyticase treatment. Spheroplasts were washed with 40 ml 1 M Sorbitol and resuspended in 5 ml/g cells Ficoll Solution (18% Ficoll, 20 mM KH_2_PO_4_ pH 7.5, 1 mM MgCl2, 0.5 mM EDTA, 1x Protease inhibitors cocktail (Roche)) for cell lysis and crude nuclei were recovered by centrifugation at 30,000 rcf for 30 minutes at 4°C.

#### Chromatin digestion

Crude nuclei were resuspended in 4 ml/g of cells in digestion buffer (1 M Sorbitol, 10 mM Tris-HCl pH 7.5, 50 mM NaCl, 1 mM CaCl_2_, 5 mM MgCl_2_, 1 mM β-mercaptoethanol and 1x Protease inhibitors cocktail) and equilibrated for 10 min in a 30°C water bath. Micrococcal Nuclease (MNase) (Sigma N5386) was diluted to 0.5 U/µl in MNase dilution Buffer (50% Glycerol, 50 mM Tris-HCl pH 8, 50 µM CaCl_2_). Crude nuclei were reacted with 5 U/g cells MNase at 30°C for 45 minutes. The reaction was quenched by addition of 1/4 volume of 5x quenching buffer (40 mM Tris-HCl pH 7.5, 2.45 M NaCl, 62.5 mM EDTA) and incubated on ice for 15 minutes to release chromatin fragments into solution. Chromatin was then cleared by centrifugation at 30,000 rcf for 30 minutes. The supernatant containing chromatin fragments was then treated with 10 µg/g cells DNase-free RNase (Roche) at 30°C for 90 minutes and subsequently cleared by centrifugation at 30,000 rcf for 30 minutes. This RNase step was critical for removing large amounts of contaminating RNA, which are likely ribosomal RNAs, to yield enzymatically active nucleosomes consistently.

#### Nucleosome purification

The cleared chromatin extract was then loaded onto a Q HP HiTrap (Cytiva) recursively at 4°C for 16 hours. Negatively charged DNA and chromatin bound to the anion exchange Q HP resin. The loaded Q HP HiTrap was washed with 20 column volumes of buffer A (5 mM Tris-HCl pH 7.5, 300 mM NaCl) and nucleosomes were eluted using twelve column volume (CV) salt gradient to reach 100% of buffer B (5 mM Tris-HCl pH 7.5, 750 mM NaCl). The peak of nucleosomes consistently eluted around 630 mM NaCl. Eluted aliquots were analyzed by polyacrylamide gel electrophoresis (PAGE), and nucleosomal fractions were pooled and concentrated to approximately 500 µl using Amicon Ultra 15 with a molecular weight cutoff of 50K (Millipore). Salt concentration was then reduced by addition of 10 volumes of 5 mM Tris-HCl pH 7.5, 50 mM NaCl, 20% Glycerol followed by further concentration. Dilutions and concentrations steps were repeated 5 times, and the sample was brought to a final volume of approximately 250 µl. The resulting nucleosomal preparation was aliquoted immediately and stored at -80°C. The flow through of the final concentration step was used as “buffer” control in subsequent experiments.

### DNA extraction

50 µl of chromatin fractions or 20 µl of final nucleosomal sample were brought to 100 µl with ddH_2_O. 1 µg DNase-free RNase (Roche) was added and reacted for 30 minutes at 30°C. Proteins were then degraded by addition of 50 µg Proteinase K (Roche) and incubation at 56°C for 90 minutes, followed by DNA purification using a QIAquick PCR purification kit (Qiagen).

### Copper reduction assay

5 µl of nucleosomes, corresponding to approximately 2-5 µg histones, or control buffer were diluted to approximal 0.2-0.5 µM in a final reaction mixture containing 67 μM Tris-HCl pH 7.5, 50 mM KOAc, 0.5 mM bicinchoninic acid (BCA) (Sigma) and 100 μM TCEP (Sigma) pH 7.5, as indicated. Zinc was added in the form of ZnSO_4_ where indicated. Magnesium, calcium, nickel, manganese or cadmium was added as MgCl_2_, CaCl_2,_ NiSO_4,_ MnCl_2_ or CdCl_2_ where indicated. Reactions were then started by addition of a CuSO_4_-Tricine-HCl pH 7 1:8 or a CuSO_4_-Histidine 1:1 mix, where indicated to a final concentration of 0.5 mM. Absorbance at 562 nm was measured every 0.5 seconds using a Hewlett-Packard HP8453 diode-array UV/Visible spectrophotometer. Initial velocities (*v*_0_) were obtained as the Ordinary Least Squares (OLS) slope of Cu^1+^ formation over the first 15 seconds. For reactions in absence of zinc no linear signal was able to fit the first 15 seconds and maximal velocity (*V*_max_) was determined as the highest slope of any sliding 60-second window. P values were determined using Welch’s two-sample *t*-test. Dose dependence of increasing zinc was tested by orthogonal polynomial contrasts on the actual zinc concentrations. Hill models were compared by extra-sum-of-squares *F*-test and Akaike information criterion (AICc). Calculations were performed using Claude Science.

### Purification of recombinant Lyticase

A plasmid for the expression of recombinant Lyticase was kindly provided by Craig Peterson (pCP330). The plasmid was transformed into BL21(DE3) (Invitrogen) and 35 ml starter culture with 2xTY containing 100 µg/ml carbenicillin (Goldbio) was grown for 16 hours. A 1:100 dilution into 3 L of the same media was grown to OD_600_ ∼0.25 and induced with 0.4 mM Isopropyl β–D-1-thiogalactopyranoside (IPTG; Goldbio). Cells were harvested, washed once with 500 ml 25 mM Tris-HCl pH 7.5 and finally stirred for 20 min in 80 ml 12.5 mM Tris-HCl pH 7.5, 1 mM EDTA, 20% sucrose and harvested again at room temperature. Cells were then resuspended in ice-cold 0.5 mM MgSO_4_ and stirred on ice for 20 min. Cell debris was then pelleted at 4°C and the supernatant was dialyzed three times against 2 L of 50 mM NaOAc pH 5.3, 1 mM EDTA, and 1 mM β-mercaptoethanol at 4°C. Precipitate was removed by centrifugation and the supernatant containing Lyticase was aliquoted and flash frozen at -80°C. Activity of the enzyme was determined empirically.

### Histone purification

Histone purification was performed as previously described (44) except that purification by size exclusion was omitted and the solubilized inclusion bodies were pre-cleaned by passage over a Hi-Trap Q-Sepharose column (GE Healthcare) before loading onto a Hi-Trap SP-Sepharose column (GE Healthcare). Histones were eluted by a salt gradient from 200 to 600 mM NaCl, as described previously (44), and fractions containing pure histones were pooled, dialyzed three times against 4 L of 2 mM β-mercaptoethanol (Sigma) in water, flash frozen and lyophilized.

### Histone H3–H4 tetramer assembly

H3–H4 tetramer assembly was performed as previously described (45) with some modifications. Briefly, equimolar amounts of H3 and H4 were dissolved in 7 M Guanidinium HCl, 20 mM Tris-HCl pH 7.5 and 10 mM DTT at a concentration of 1-2 mg/mL total protein. refolding was achieved by 1-2 hr dialysis at 4°C against 1 L of 2 M NaCl, 10 mM Tris-HCl pH 7.5, 1 mM EDTA, 5 mM β-mercaptoethanol, then 2 hrs against 1 L of 1 M NaCl, 10 mM Tris-HCl pH 7.5, 1 mM EDTA, 5 mM β-mercaptoethanol, followed by 16 hrs against 0.5 M NaCl, 10 mM Tris-HCl pH 7.5, 1 mM EDTA, 5 mM β-mercaptoethanol. The refolded tetramer was centrifuged for 5 min at 13000 rcf at 4°C to eliminate insoluble particles and purified by size exclusion chromatography on a HiLoad 16/600 Superdex 200 column (Cytiva) at 1 mL/min in 500 mM NaCl, 5 mM Tris-HCl pH 7.5. Fractions containing the tetramer were concentrated using Amicon Ultra-15/Ultracel-30K centrifugal filters (Millipore). The flow-through buffer was used as negative control (“buffer”) in the copper reductase assay.

### Mass Spectrometry

Protein bands were excised from the gel and in-gel digested with trypsin according to a published protocol (46), with some modifications. A piece of gel cut from a blank region of the gel and processed in parallel with the sample, served as a control. Briefly, the gel pieces were destained, dehydrated and dried. The proteins in the gel pieces were reduced with 10 mM dithiothreitol at 56°C followed by alkylation with 55 mM iodoacetamide at 25°C in dark. After washing, the gel pieces were subjected to dehydration by addition of acetonitrile, then rehydration in 50 mM NH_4_HCO_3_ followed by dehydration again. The gel pieces were dried and rehydrated in 12.5 ng/μL of trypsin in 50 mM NH_4_HCO_3_ at 4°C. After complete rehydration of gel pieces, the supernatant was removed and replaced with 50 mM NH_4_HCO_3_ buffer (without trypsin) and incubated at 37°C, overnight. Peptides were extracted from the gel matrix by two rounds of incubation in extraction buffer containing 5% formic acid and 30% acetonitrile. Following extraction, the gel pieces were dehydrated with acetonitrile. The supernatant was pooled with extraction buffer recovered from the previous step, dried and subsequently analyzed by LC-MS/MS.

Briefly, peptides were separated by reversed phase chromatography using 75 μm inner diameter fritted fused silica capillary column packed in-house to a length of 25 cm with bulk 1.9 µM ReproSil-Pur beads with 120 Å pores (47). The increasing gradient of acetonitrile was delivered by a Dionex Ultimate 3000 (Thermo Scientific) at a flow rate of 200 nl/min. The MS/MS spectra were collected using data dependent acquisition on Orbitrap Fusion Lumos Tribrid mass spectrometer (Thermo Fisher Scientific) with an MS1 resolution (r) of 120,000 followed by sequential MS2 scans at a resolution (r) of 15,000. The data generated by LC-MS/MS were analyzed on MaxQuant bioinformatic pipeline (48). The Andromeda integrated in MaxQuant was employed as the peptide search engine and the data were searched against *Saccharomyces cerevisiae* database (Uniprot Reference UP000002311). A maximum of two missed cleavages was allowed. The maximum false discovery rate for peptide and protein was specified as 0.01. Label-free quantification (LFQ) was enabled with LFQ minimum ratio count of 1. The parent and peptide ion search tolerances were set as 20 and 4.5 ppm respectively.

### Inductively-coupled plasma mass spectrometry (ICP-MS)

Sample preparation and analysis was performed as described in (3).

### Flow cytometry

Single colonies of cells bearing the Cup2 reporter plasmid (3) were grown in liquid SC-ura media containing 0.25 µM ZnSO_4_ (“low zinc”) for 16 hrs to deplete endogenous zinc reserves. Cells are then washed twice with ddH_2_O and seeded at OD_600_ 0.2 density in either low zinc SC-ura or low zinc SCEG-ura and 0, 1 or 10 µM CuSO_4_ was added. After 24 hrs of growth, cells were directly assayed in the liquid media using a BD LSRFortessa^TM^ X-20 instrument. Fluorescine Isothiocyanate (FITC) signal was collected using a 488 nm laser for excitation and a 515 – 545 nm bandpass filter for emission. About 50,000 events were collected on the cytometer and gated by forward and side scatter parameters to analyze single cells.

### Yeast growth curves

Yeast was inoculated from plate for a pre-culture in SC or SCEG without Zinc, in order to deplete intracellular zinc reserves. After 24 hrs, cells were diluted to OD_600_ in the same media with the addition of 0.02, 0.1 or 0.25 µM ZnSO_4_ (low zinc media) and, where indicated, additional CuSO_4_. 2.2 µM ZnSO_4_ was added for all replete zinc growth conditions. Optical density was determined at indicated times using a SmartSpec 3000 (Biorad) spectrophotometer. Results were curve fit using the modified Gompertz model as described in Zwietering *et al.*, 1990 (49) and lag phase and doubling time were derived accordingly. The significance of growth differences between curves was determined using the *F*-test, while lag phase and doubling time were compared with Welch’s two-sample *t*-test assuming unequal variance. Calculations were performed using Claude Science. Curve fitting used scipy.optimize.curve_fit.

## Acknowledgments

This study was supported by the NIH grants GM158109 and GM140106 and Moore-Simons Project on the Origin of the Eukaryotic Cell to SKK. Flow cytometry was performed in the UCLA Jonsson Comprehensive Cancer Center (JCCC) and Center for AIDS Research Flow Cytometry Core Facility that is supported by National Institutes of Health awards P30 CA016042 and 5P30 AI028697, and by the JCCC, the UCLA AIDS Institute, the David Geffen School of Medicine at UCLA, the UCLA Chancellor’s Office, and the UCLA Vice Chancellor’s Office of Research. We thank Craig L. Peterson for kindly providing the Lyticase expression plasmid pCP330, and all members of the Kurdistani lab for helpful discussions. We also thank Vijaya Pandey and James Wohlschlegel of the UCLA Proteome Research Center for mass spectrometry analyses and Stefan Schmollinger and Sabeeha Merchant for help with ICP-MS.

## Data availability

The data underlying this article are available in the article and in its online supplementary material.

## Supplement

**Supplement Figure S1.**
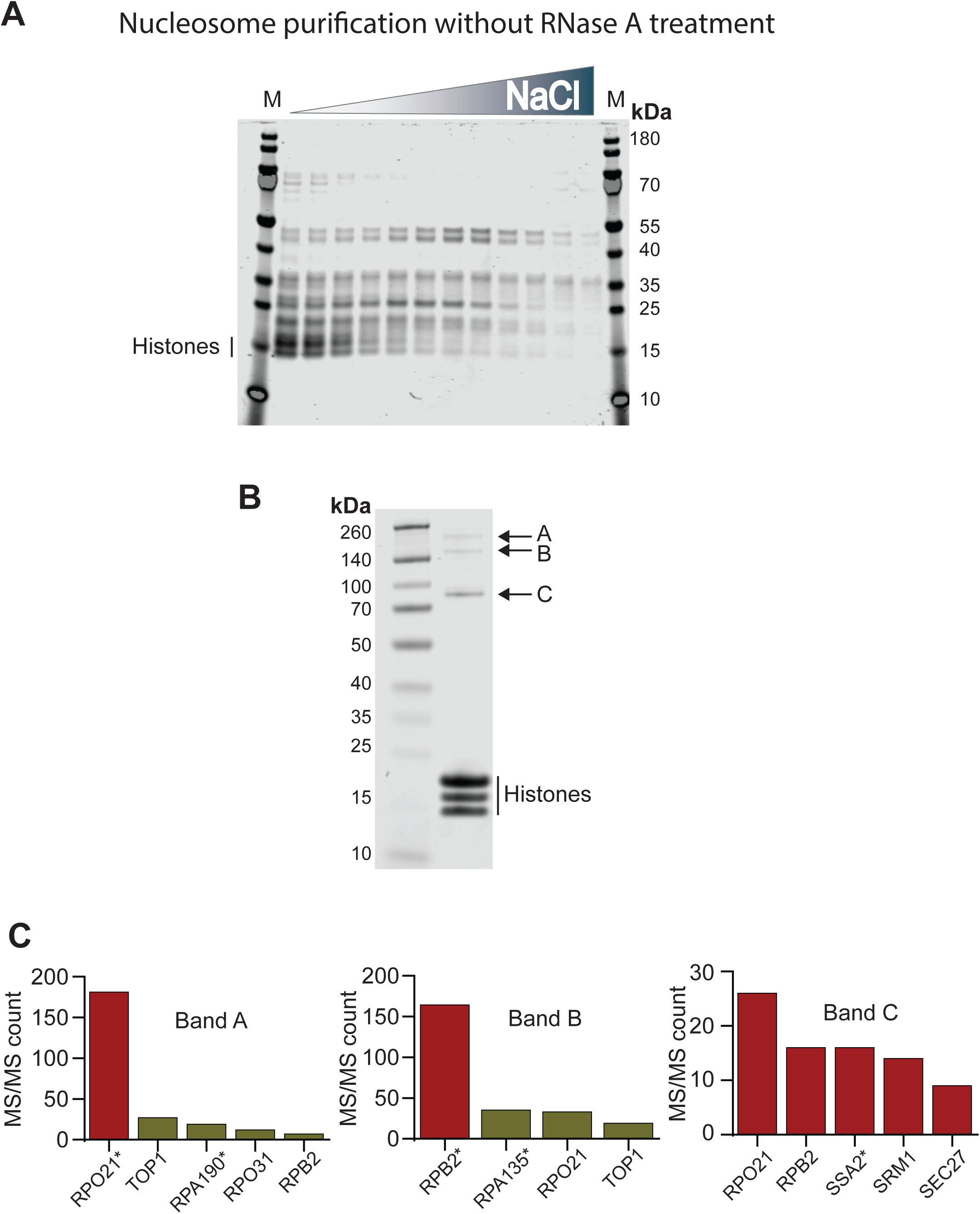
Co-eluting proteins in nucleosome preparations with and without RNase A treatment. A, Coomassie-stained PAGE of eluted nucleosomal fractions prepared as in Fig. 1C but without the RNase A treatment step. B, PAGE of a representative purified nucleosome preparation. The major co-eluting protein bands are labeled A, B, and C. C, Bar graphs showing normalized MS/MS counts for proteins identified by mass spectrometry in each co-eluting band shown in (B). Proteins whose predicted molecular weights are consistent with the corresponding gel band are indicated by an asterisk.

**Supplement Figure S2.**
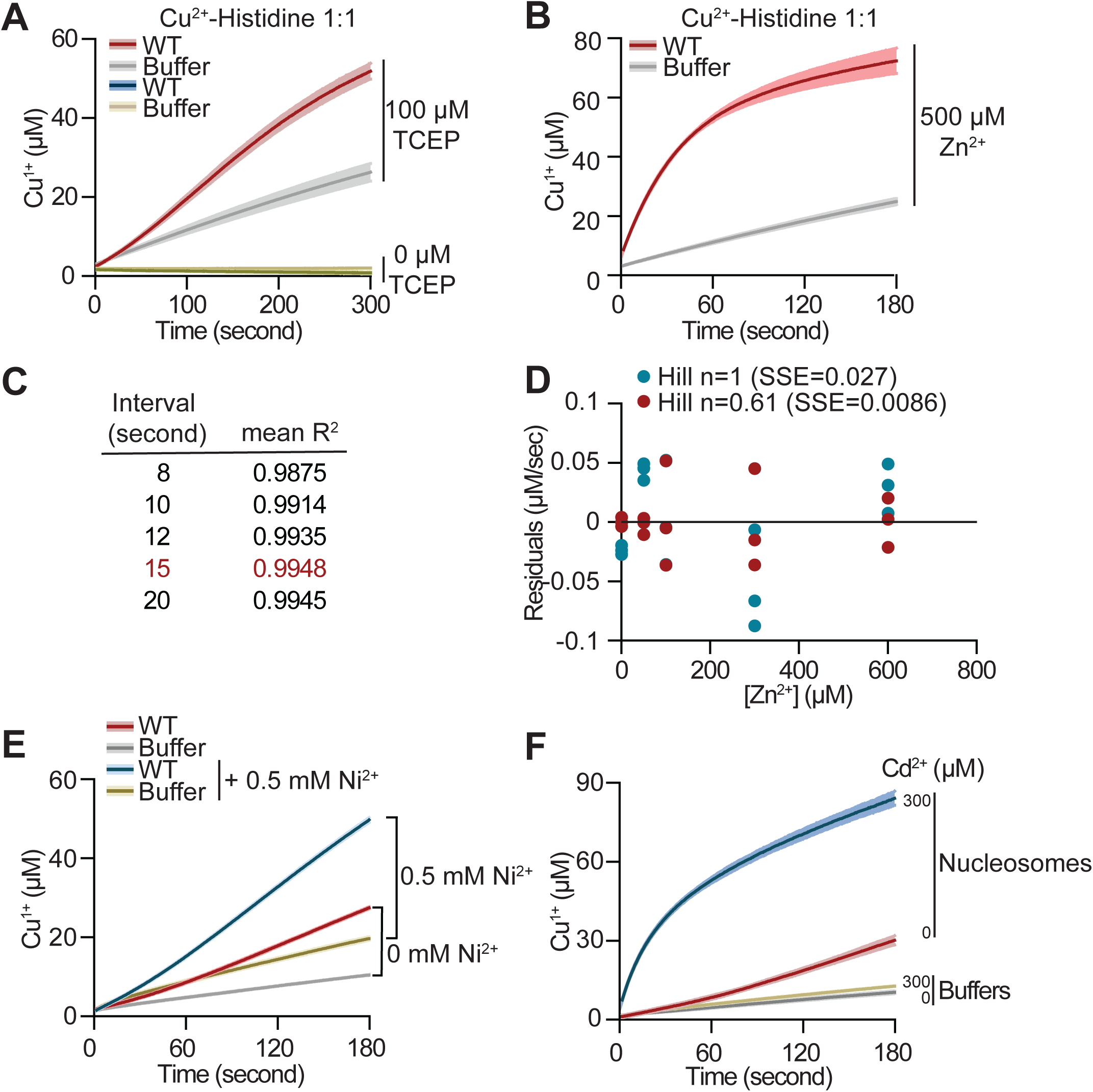
Zinc selectively enhances nucleosome copper reductase activity. A, Progress curves of a Cu^2+^ reduction by WT nucleosomes using a 1:1 Cu^2+^-histidine complex as substrate in presence or absence of 100 µM TCEP. B, Progress curves of a Cu^2+^ reduction in presence of the indicated amount of Zn^2+^ using a 1:1 Cu^2+^-histidine complex as substrate. C, Determination of the time interval used to estimate initial reaction velocity. Mean R^2^ values were calculated for linear fits over the indicated intervals; the 15-s interval yielded the mean R^2^ value closest to 1. D, Residuals for the Zn^2+^ dose-response data shown in Fig. 2C relative to Hill-model fits with n fixed at 1 or allowed to vary (n = 0.61). The free-Hill fit (n = 0.61) produced a lower sum of squared errors (SSE = 0.0086) than the fit with n = 1 (SSE = 0.027), indicating a better fit to the data. E, Progress curves of Cu^2+^ reduction by WT nucleosomes or buffer control in the presence or absence of Ni^2+^. Nickel primarily enhanced non-specific Cu^2+^ reduction, as reflected by shifts in both buffer and nucleosome curves. Lines and shading represent the mean ± SD of two or three assays. F, Progress curves of Cu^2+^ reduction by WT nucleosomes in the presence of indicated Cd^2+^ concentrations. Lines and shading represent the mean ± SD of three assays.

**Supplement Figure S3.**
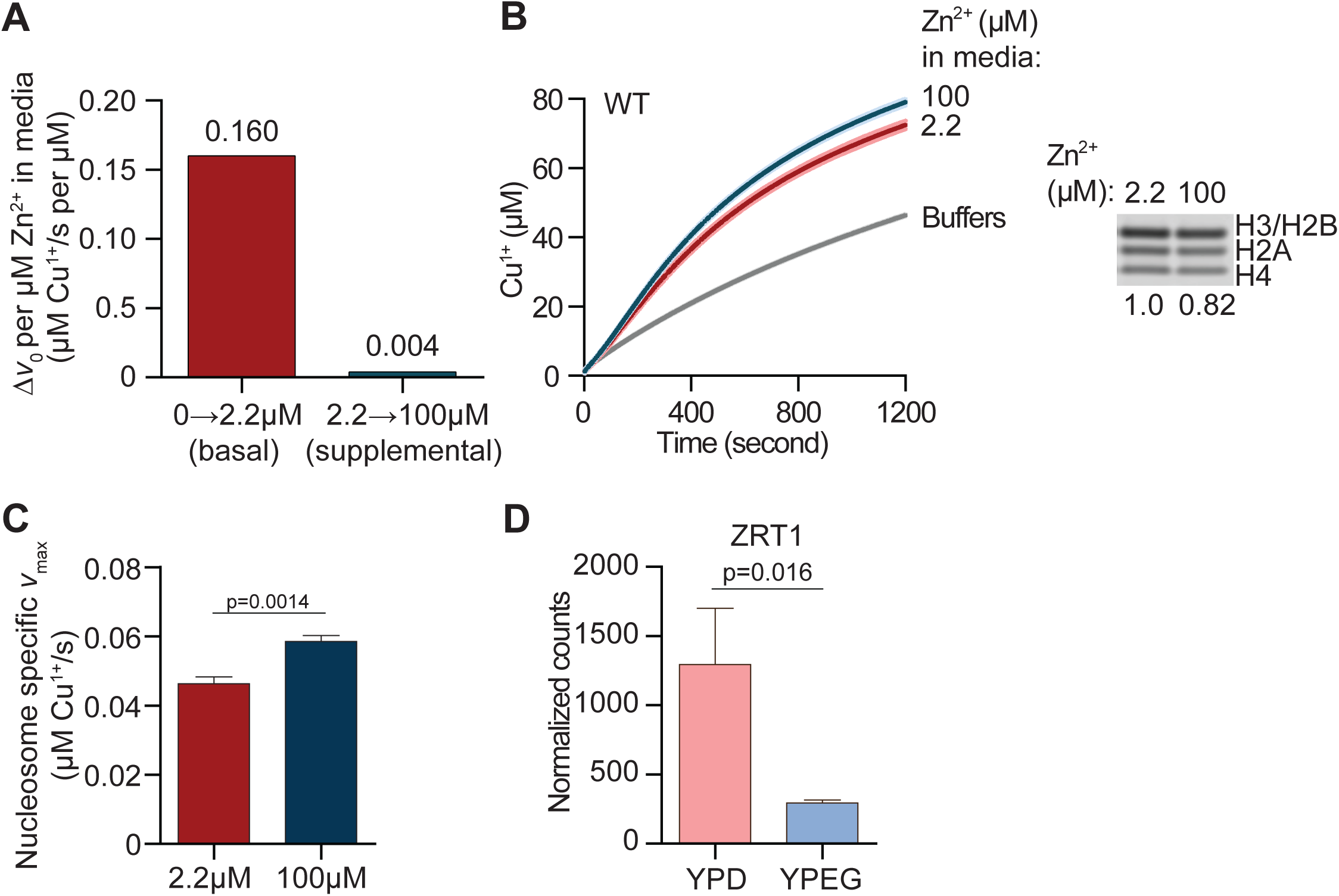
Availability of zinc in growth media determines copper reductase activity of nucleosomes. A, Increase in nucleosome-specific initial velocity (Δ*v*₀) per µM Zn^2+^ added to the growth medium, calculated for the transition from zinc-free to basal zinc conditions (0 to 2.2 µM) and from basal to zinc-supplemented conditions (2.2 to 100 µM). B, Left, progress curves of Cu^2+^ reduction by WT nucleosomes purified from cells grown in medium containing the indicated concentrations of Zn^2+^ and assayed without added ZnSO_4_. Right, Coomassie-stained gel of nucleosomes used in the assay, with relative loading normalized to nucleosomes isolated from cells grown in 2.2 µM Zn^2+^. C, Nucleosome-specific maximal reaction velocities (*V*_max_) calculated from the progress curves shown in(B). D, *ZRT1* expression, shown as normalized mRNA-seq counts, in WT cells grown under fermentative (YPD) or respiratory (YPEG) conditions. P values were calculated using Welch’s *t*-test.

**Supplement Figure S4.**
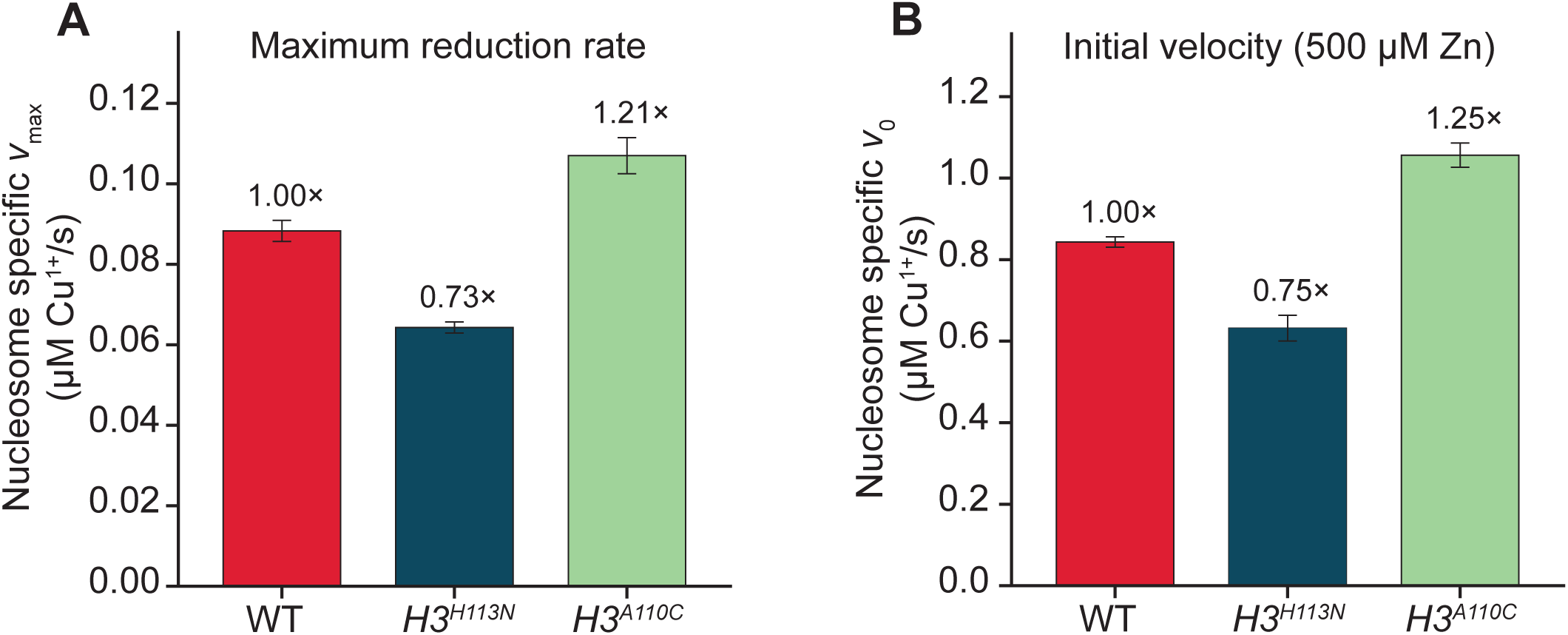
Mutations at the histone H3 catalytic site decrease or enhance native nucleosome copper reductase activity independently of zinc addition to the assay. A, Maximum velocities (*V*_max_) of Cu^2+^ reduction by WT, H3H113N and H3A110C nucleosomes without Zn^2+^ added to the *in vitro* reaction. Rates relative to WT are indicated. B, Initial velocities (*v*_0_) by WT, H3H113N and H3A110C nucleosomes with 500 µM Zn^2+^ added to the *in vitro* reaction. Rates relative to WT are indicated.

**Supplement Figure S6.**
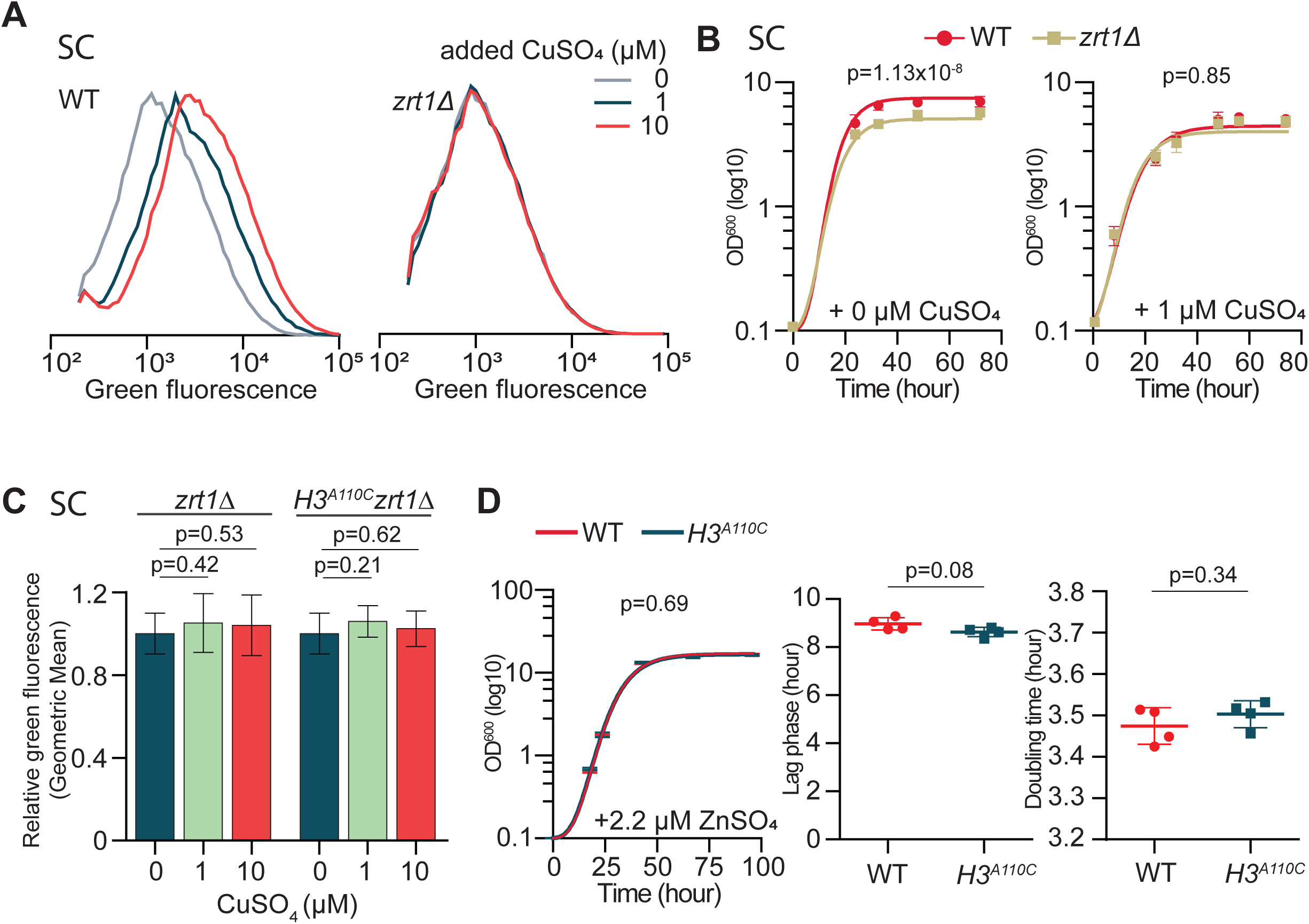
Zinc availability in growth medium influences histone H3-dependent Cu^1+^ bioavailability and growth in cells. A, Average green fluorescence distributions of WT or *zrt1Δ* cells containing the p^(CUP1)^-GFP plasmid cultured in fermentative media lacking uracil (SC-ura) containing 0.25 µM ZnSO_4_, in the absence (0) or presence of either 1 or 10 µM additional CuSO_4_ from eight experiments. B, Growth curves in fermentative SC media containing 0.25 µM ZnSO_4_, in the absence (0; left panel) or presence 1 µM additional CuSO4. Lines show means at each time point ± SD from three experiments. C, Average relative geometric means of green fluorescence *zrt1Δ* or *H3^A110C^zrt1Δ* cells containing the pCUP1-GFP plasmid cultured in SC-ura containing 0.25 µM ZnSO_4_, in the absence (0) or presence of either 1 or 10 µM additional CuSO_4_ from eight experiments. D, Growth after the indicated hours in SCEG in replete zinc concentrations (2.2 µM ZnSO_4_) (left). Graphic representation of the time of lag-phase (middle) and doubling time (right) are reported. P values were calculated using the *F*-test for growth curves and a Welch’s *t*-test for bar graphs.

**Supplement Figure S6.**
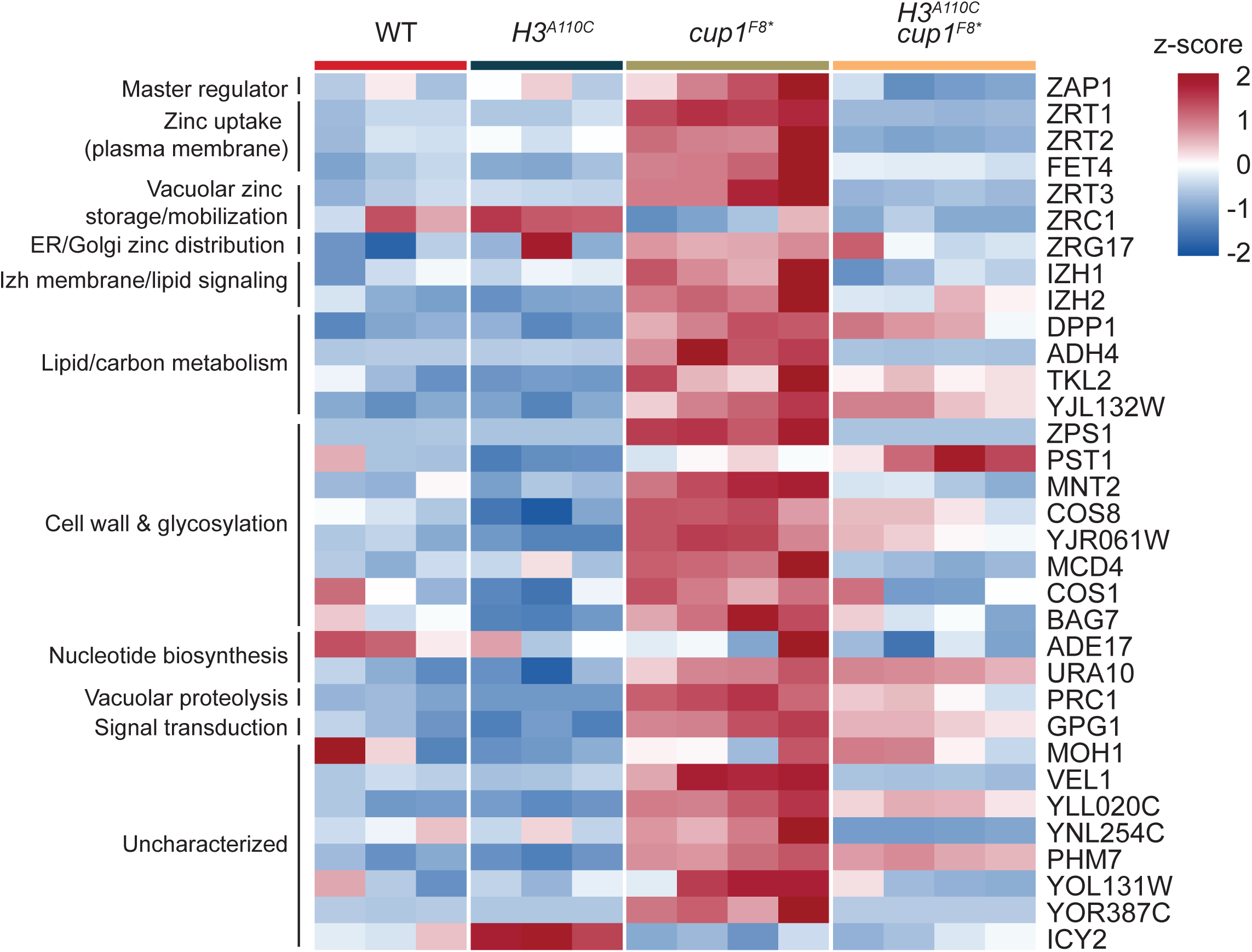
H3A110C suppresses transcriptional changes in zinc-related genes associated with loss of the Cup1 metallothionein. Heat map of expression levels of genes involved in zinc homeostasis or zinc-related processes in WT, *H3^A110C^*, *cup1F8\**, and *H3^A110C^cup1F8\** cells. Genes were selected based on the zinc-related gene set described by Wu *et al.* in table 1 (50), and expression values were obtained from RNA-seq data reported by Cheng *et al.* (13). Genes are grouped according to functional annotation. Each column represents data from one experimental replicate. Expression values are displayed as z-scores, with higher and lower relative expression indicated by positive and negative z-scores, respectively. The *cup1F8\** mutation is associated with broad induction of zinc-homeostasis genes, including *ZAP1*, *ZRT1*, *ZRT2*, *FET4*, and *ZRT3*, whereas introduction of H3A110C largely shifts their expression toward the WT pattern.

**Table S1:** Yeast strains used in this study.

| Strain name | Mutant short name | Description | Reference |
| --- | --- | --- | --- |
| **OCY1131** | WT | *MATa his3Δ1 leu2Δ0 met15Δ0 ura3Δ0, (hht2Δ::ura3)Δ::HHT2* | Attar et al., 2020 |
| **OCY1136** | H3<sup>H113N</sup> | *MATa his3Δ1 leu2Δ0 met15Δ0 ura3Δ0, (hht2Δ::ura3)Δ::HHT2-H113N HHT1-H113N* | Attar et al., 2020 |
| **NAY616** | H3<sup>A110C</sup> | *MATa his3Δ1 leu2Δ0 met15Δ0 ura3Δ0, (hht2Δ::ura3)Δ::HHT2, hht2-A110C, hht1-A110C* | Attar et al., 2020 |
| **OCY1981** | cup1<sup>F8stop</sup> | *MATa his3Δ1 leu2Δ0 met15Δ0 ura3Δ0, (hht2Δ::ura3)Δ::HHT2, cup1-F8stop* | Attar et al., 2020 |
| **OCY1984** | H3<sup>A110C</sup>cup1<sup>F8*</sup> | *MATa his3Δ1 leu2Δ0 met15Δ0 ura3Δ0, (hht2Δ::ura3)Δ::HHT2, hht2-A110C, hht1-A110C, cup1-F8stop* | Attar et al., 2020 |
| **AKY84** | zrt1Δ | *MATa his3Δ1 leu2Δ0 met15Δ0 ura3Δ0, (hht2Δ::ura3)Δ::HHT2, zrt1Δ* | This study |
| **CCY0402** | H3<sup>A110C</sup> zrt1Δ | *MATa his3Δ1 leu2Δ0 met15Δ0 ura3Δ0, (hht2Δ::ura3)Δ::HHT2, hht2-A110C, hht1-A110C, zrt1Δ* | This study |

## Notes

### Competing Interest Statement

The authors have declared no competing interest.

### Summary of Updates

An entirely new set of data strengthens the conclusion that zinc regulates copper reduction by the yeast nucleosome. Importantly, we now show that the in vivo availability of zinc establishes a catalytically competent nucleosome state during cell growth.

